# Microfluidic device for on-chip mixing and encapsulation of lysates

**DOI:** 10.1101/247627

**Authors:** Chang Jui-Chia, Swank Zoe, Keiser Oliver, Maerkl Sebastian, Amstad Esther

## Abstract

Emulsion drops are often employed as picoliter-sized containers to perform screening assays. These assays usually entail the formation of drops encompassing discrete objects such as cells or microparticles and reagents to study interactions between the different encapsulants. Drops are also used to screen influences of reagent concentrations on the final product. However, these latter assays are less frequently performed because it is difficult to change the reagent concentration over a wide range with high precision within a single experiment. In this paper, we present a microfluidic double emulsion drop maker containing pneumatic valves that enable injection of different reagents using pulsed width modulation and subsequent mixing. This device can produce drops from reagent volumes as low as 10 μl with minimal sample loss, thereby enabling experiments that would be prohibitively expensive using droplet generators that do not contain valves. We employ this device to monitor the kinetics of cell free synthesis of green fluorescent proteins inside double emulsions. To demonstrate the potential of this device, we perform DNA titration experiments in double emulsion drops to test the influence of the DNA concentration on the amount of green fluorescence proteins produced.

## Introduction

Emulsion drops are well-suited containers for performing chemical and biochemical reactions under well-defined conditions and in volumes that are significantly smaller than those required to conduct reactions in bulk. This is especially beneficial for high throughput screening assays.^1–3^ The accuracy of such assays depends on the degree of control over the composition and concentration of reagents contained inside the drops as well as their size distribution. Drops with a narrow size distribution can be produced using microfluidics.^4–6^ These drops have, for example, been employed as containers for drug screening assays,^7, 8^ to perform polymerase chain reactions (PCR) from viruses,^9, 10^ or single cells,^11^ for directed evolution of enzymes,^12^ to study the secretion of proteins on a single cell level,^13^ or to identify genes that are responsible for a certain cellular phenotype.^14^ To perform these screening assays, reagents are often pre-mixed before they are injected into the device. Pre-mixing limits kinetic studies to characterizing very slow reactions or the late stages of faster reactions because reagents start to react before they are loaded into drops. Moreover, pre-mixing prevents *in situ* changes of the relative reagent concentrations such that only one solution composition can be screened per experiment. A possibility to overcome these shortcomings is the injection of reagents into drops after they have been formed for example through the application of high electric fields^15–19^ or the addition of chemicals that destabilize drops.^20^ However, the number of different reagents that can be controllably added to intact drops is limited. Moreover, it is difficult to accurately and continuously vary the concentration of injected reagents. A possibility to gradually and controllably vary the reagent concentration is to co-flow two fluids under laminar conditions; in this case, the reagent exchange is diffusion limited.^21–23^ However, because mixing relies on diffusion, the spatio-temporal control over the solution composition is poor. This control can be improved if mixing is enhanced, for example by introducing turbulences into the fluid flow using structured microchannels^24, 25^ or active mixers, such as micropumps, or micromixers.^26^ However, it always takes some time to equilibrate injection flow rates especially if multiple fluids are involved. Thus, even with mixing features being implemented, it is difficult to controllably and continuously change the concentrations of different reagents with high temporal resolution.

The concentration of reagents can be changed over a wide range and on very short time scales if microfluidic channels are equipped with pneumatic valves that can be opened and closed rapidly. These valves allow the formation of a train of alternating plugs of different types of miscible liquids that can be subsequently mixed. The length of the plug of each fluid scales with the duty cycle, corresponding to the fraction of time one valve is opened compared to the entire cycle time, and can be adjusted *in situ*. This procedure enables varying relative reagent concentrations over a wide range within a single experiment by gradually changing the duty cycles; thereby only minimal volumes of reagents are consumed. This method, the so-called pulsed width modulation (PWM), has been implemented in microfluidic devices^27, 28^ and is often employed to synthesize biopolymers, to study the influence of their composition on their function, and to test the effect of certain molecules on the cell behavior.^29–34^ Recent advancements of this technology enable independent injection of up to six different reagents and changing their concentrations *in situ* by up to five orders of magnitudes.^35^ This level of compositional control over such a wide concentration range is difficult to achieve with co-flowing fluids. Despite these distinct advantages of PWM, on-chip mixing of solutions that are subsequently processed into drops of defined sizes is thus far done through co-flow of different fluids. The ability to rapidly and controllably change the concentration of reagents over a much wider range would open up new possibilities for high throughput screening assays. However, pneumatic valves have never been implemented into microfluidic flow focusing drop makers and consequently PWM has thus far not been performed in these devices.

In this paper, we present a microfluidic flow focusing drop maker that has three inlets for reagents, each of them controlled by a pneumatic valve. This device allows separate injection of different reagents using PWM, on-chip mixing, and formation of double emulsion drops of defined dimensions using this mixture as an inner phase. The pneumatic valves provide an additional benefit: they enable encapsulation of liquids with volumes as low as 10 μl at an efficiency approaching 100%. This device is employed to encapsulate lysates that synthesize green fluorescent protein (GFP) inside double emulsion drops. We demonstrate the potential of the device to perform characterization assays by titrating DNA and measuring the influence of its concentration on the amount of GFP produced in double emulsion drops.

## Results

We fabricate microfluidic devices using soft lithography.^36^ These devices contain three inlets for aqueous solutions that form the inner phase, one inlet for the oil that constitutes the middle phase, and one inlet for an aqueous phase that forms the surrounding liquid. To control the fluid flow in the three inlets for the innermost phase, we introduce pneumatic valves on top of these channels, as schematically shown by the blue features in Figure 1a.^36^ The valves enable injection of different fluids using pulsed width modulation, as exemplified in the optical micrograph in Figure 1b.^35, 37, 38^ To accelerate the mixing of the different injected reagents, herringbones are introduced to the serpentine-like section of the main channel located between the inlets for the innermost phases and that for the oil phase,^25^ as schematically shown in Figure 1a. To separate reagent-loaded drops from empty ones, the device has a T-junction used as a sorting unit, as shown schematically in Figure 1a and in the optical micrograph in Figure 1c. The flow of double emulsions is again controlled with pneumatic valves that change the hydrodynamic resistance of the outlet channels.

**Figure 1:**
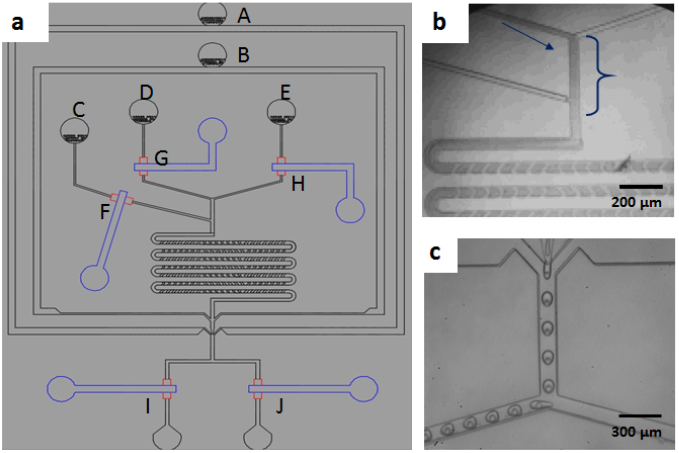
Schematic illustration of the microfluidic device (a) Overview of the device with the inlet for (A) the outermost aqueous phase, (B) the oil phase and (C-E) the three innermost aqueous phases. Each inlet for the innermost phase contains a pneumatic valve that enables controlling the fluid flow (F-H); the control lines are indicated in blue. Drops exit the device through one of the two outlet channels that also contain pneumatic valves (I-J). (b) Optical micrograph of an operating device where an aqueous phase and a dye are alternatively injected using PWM. (c) Optical micrograph of double emulsions that are collected through the left outlet.

To precisely tune the amount of liquid injected into the main channel, the switching times of the valves must be accurately controlled. Valves close if their control channels are pressurized such that the valves are pressed towards the bottom of the fluid channel. We employ solenoid valves to pressurize and depressurize the control channels; these valves are electronically controlled. To determine the response time of the valves, we use a colored fluid and quantify the intensity profile across the fluid channel underneath the valve as a function of the pulse width. If the valve is open, the entire inlet channel underneath the valve is colored, as shown in the optical micrograph in Figure 2a; in our device, the inlet channel section below the valve is 200 μm wide. If the valve is closed, the fluid is pushed aside such that this section becomes transparent, as shown in Figure 2b. If the pulse width of the valve, corresponding to the time the valve is open, is 20 ms, no fluid can pass, as demonstrated by the flat curve of the black circles in Figure 2c. If we increase the pulse width to 30 ms, some fluid passes and even more fluid passes if the pulse width is increased to 40 ms, as indicated by the increased peak intensity, shown by the blue squares and red stars in Figure 2c. Increasing the pulse width further broadens the intensity peak but does not increase its amplitude any more, as shown in Figure 2c. These results indicate that the response time of the valves, and thus their minimum pulse width, is 40 ms. This pulse width sets a lower limit to the size of plugs that can be formed. This size depends on the fluid flow rate that we set to 250 μl/h; in this case, the minimum plug volume is 3.5 nl.

**Figure 2:**
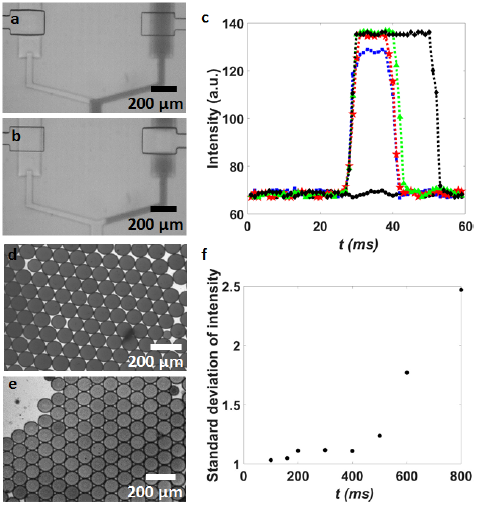
Characterization of the pneumatic valves. (a, b) Optical micrographs of a pneumatic valve that is (a) opened and (b) closed. (c) The intensity of the colorant measured across the channel underneath the valve for pulse widths of 10 ms, (●) for 20 ms; (ƞ) for 25 ms; (Ȍ) for 30 ms; (ƴ) for 40 ms; (ʆ) for 50 ms. (d, e) Optical micrographs of drops formed with a pulse width of (c) 100 ms and (d) 800 ms. (f) The standard deviation of the dye intensity in drops as a function of the pulse width.

A key feature of the device is its ability to precisely change *in situ* the composition of reagents that are encapsulated in double emulsions. This degree of control is only possible to obtain, if adjacent plugs containing different reagents are homogeneously mixed before the solution is compartmentalized into drops. The longer the plugs are, the longer it will take to convert an array of plugs into a homogeneous solution because the diffusion lengths are increased. To quantify the maximum length of plugs that our device can mix before the solution is broken into drops, we employ two liquids, water and a water-soluble dye, as an inner phase. The lengths of the plugs are varied by changing the pulse width of the two corresponding valves. When the aqueous phase reaches the water-oil junction, it is broken into 70 μm diameter drops. If the two aqueous phases are fully mixed when the solution reaches the water-oil junction, the intensity of the drops is homogeneous, as indicated in Figure 2d. By contrast, if the plugs are too long, the composition of the solution at the oil-water junction varies over time and the intensity of the drops is heterogeneous, as shown in Figure 2e. To quantify the maximum length of aqueous plugs that can be fully mixed in our device, we measure the standard deviation of the drop intensity as a function of the pulse width; these experiments are performed using liquids whose viscosity is close to that of water. If the pulse width is below 400 ms, the intensity of drops is uniform, as shown in the optical micrograph in Figure 2d and summarized in Figure 2f. By contrast, if the pulse width is increased above this value, the intensity becomes heterogeneous, as shown in the optical micrograph in Figure 2e and summarized in Figure 2f. These results indicate that the maximum pulse width is 400 ms corresponding to a plug volume of 35 nl. Hence, in our device, we can vary the plug volumes from 3.5 to 35 nl. However, these are not fundamental limits. The dynamic range of the device could be increased by prolonging the mixing unit, or by altering the channel dimensions.

Microfluidics allows encapsulation of reagents with a high efficiency. However, in most cases, some reagents are lost during device start-up. This fluid loss poses no problem if reagents are inexpensive and available in large quantities. However, some biological assays involve expensive reagents or samples that are only available in very small volumes. To process smaller volumes in microfluidic devices, sample losses must be minimized. This can be achieved using pneumatic valves because they allow initialization of the device with water prior to switching to the expensive reagent. When the device runs stably, the channel for the water phase is closed while channels containing expensive reagents are opened. To separate reagent-loaded drops from empty ones, we again employ pneumatic valves that are incorporated into the collection channels. To test the performance of these valves, we form aqueous single emulsion drops and collect them through the right outlet. Even if the right valve is open, 80 μm diameter drops tend to break when they pass it because the channel height underneath the valve cannot exceed 14 μm for them to be able to fully close the channel. To prevent drop break-up, we increase the height underneath the sorting valves to 20 μm. These valves do not completely close the channel even if they are fully pressurized. Instead, they decrease the height of the channel, thereby increasing its hydrodynamic resistant to such an extent that under our operating conditions drops do not pass this obstacle any more. We exemplify this behavior by closing the valve of the left collection channel and opening the one of the right channel. In this case, drops do not break when they pass the open valve and can be collected on the right side, as indicated in the optical micrograph in Figure S1a. To maximize the accuracy of the sorting, we close the valve of the right channel 10 ms before we open the valve of the left channel: This procedure reduces pressure differences between the two collection channels and therefore allows the drops to flow into the desired collection channel as soon as the corresponding valve is opened. If the injection rate is below 600 μl/h, drops can be sorted without any loss, as shown by the optical time-lapse images in Figure S1a. By contrast, if this injection rate is exceeded, drops tend to split at the sorting junction during the switching operation, as shown in the optical time laps images in Figure S1b.

For drops to be used as reaction vessels, they must be stable during incubation. To test the stability of drops, we load them with lysates^39, 40^ and a solution containing 30 mM 3-PGA and incubate them at 29°C for 3 h. Unfortunately, most of the single emulsion drops coalesce during their incubation such that the resulting sample is polydisperse. This coalescence limits the usability of drops for many screening applications. The stability of drops coated with perfluorinated triblock copolymers can be compromised by the presence of high concentrations of ions.^41^ To test if we can increase the drop stability, we decrease the concentration of 3-PGA to 4 mM. However, also in this case, single emulsion drops tend to coalesce albeit to a smaller extent, as summarized in Figure 3a and shown in the fluorescence micrographs in Figures S2a and S2b. We expect the coalescence of single emulsion drops to be caused by ions that are in close proximity to the surfactants or to drop interfaces. If this expectation holds, double emulsions should be more stable against coalescence because their outer liquid-liquid interface is separated from ions by the oil shell. Indeed, the percentage of intact double emulsions is significantly higher than that of single emulsion drops, even if a solution containing 30 mM 3-PGA is used as an innermost phase, as summarized in Figure 3a.

To test if we can express GFP in double emulsions, we employ PURE, a cell-free transcription / translation reaction mixture generated from purified components.^42^ PURE is now commercially available (NEB) but is a relatively expensive reagent; it currently costs over 1 USD per μl. Therefore, it is beneficial to only consume small volumes. Small volumes are difficult to handle with standard microfluidic devices where volumes between 50 and 100 μl can easily be lost during start-up. To prevent sample loss during start-up of our device, deionized water is injected through inlet C. When the device runs stably, valve F is closed, an aqueous solution containing PURE is injected through inlet D, and an aqueous solution containing DNA is injected through inlet E. The two reagent-containing solutions are injected using PWM with pulse widths of 50 ms. The aqueous mixture is broken into 65 μm diameter double emulsion drops that display a narrow size distribution. These drops are incubated at 29°C for 3 h and the formation kinetics of functional GFP is measured by acquiring a fluorescent micrograph every 8 min. Within 2 h fluorescence reaches a plateau as shown in the fluorescence intensity trace in Figure 3b. These experiments show feasibility to synthesize GFP in double emulsions and the economic use of expensive reagents such as PURE.

**Figure 3:**
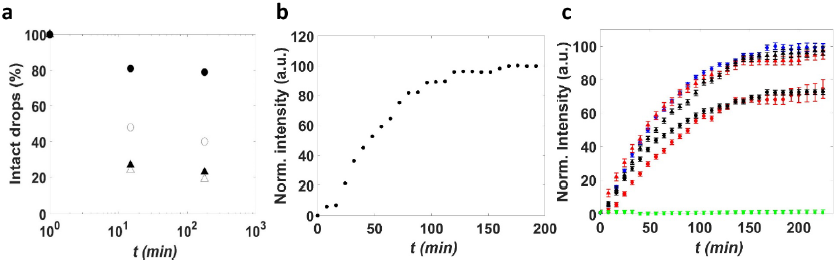
Synthesis of GFP in drops. (a) The percentage of drops that remain intact as a function of the incubation time for single emulsion drops loaded with 30 mM (Ƶ) and 4 mM 3-PGA (ƴ) and double emulsion drops containing 30 mM (○) and 4 mM 3-PGA (●). (b) The normalized fluorescence intensity measured in double emulsion drops loaded with PURE as a function of the incubation time. (c) The normalized fluorescence intensity of lysate solutions in bulk (ʆ), in single emulsion drops loaded with lysates through PWM (ƴ) and with pre-mixed lysates (ƴ), and double emulsion drops loaded with lysates through PWM (●) and with pre-mixed lysates (●) as a function of the incubation time. As a control, lysate extract was loaded into drops in the absence of DNA (ƞ).

Double emulsions are significantly more stable than single emulsions. Nevertheless, a large fraction of the cores containing 30 mM 3-PGA and lysate extracted from *E. coli* merge with the surrounding aqueous phase such that all the encapsulants are released, as shown in Figure 3a. To increase the fraction of intact double emulsion drops, we reduce the 3-PGA concentration to 4 mM. This reduction in 3-PGA concentration significantly increases the percentage of intact double emulsions, as exemplified in the fluorescence micrographs in Figures S2c, d and summarized in Figure 3a. Hence, solutions containing 4 mM 3-PGA are employed for the following experiments.

To investigate the kinetics and amount of synthesized functional GFP, we quantify the fluorescence intensity of each drop. Encapsulation of lysates into 65 μm diameter drops does not alter the kinetics of GFP synthesis, as summarized in Figure 3c. However, the amount of GFP produced in double emulsions is significantly lower than that produced in single emulsion drops and in bulk, as indicated by the blue diamond in Figures 3c. In our experiments, single emulsion drops encompass 0.55 μM functional GFP whereas double emulsion drops only contain 0.39 μM functional GFP, as summarized in Figure S3. This difference might be caused by a partial leakage of reagents through the shell of double emulsions, by analogy to the crosstalk observed between aqueous drops that are dispersed in perfluorinated oils.^43^ Nevertheless, there is a significant amount of functional GFP synthesized in double emulsion drops, indicating that they can be used to screen effects of synthesis conditions on the amount of functional GFP produced.

The main feature of our microfluidic device is its ability to change the reagent concentrations *in situ* with high accuracy. To exploit this feature, we perform DNA titration experiments: Water is injected through inlet D, an aqueous solution containing lysate extracts from *E. coli*, 30 mM 3-PGA, and an energy source through inlet C, and an aqueous solution containing 15 nM DNA through inlet E. The cycle time of the valves is kept constant at 4 s while the pulse width of valves G and H are varied from 40 ms to 320 ms in four steps. Using this procedure, we screen four DNA concentrations, 13.3, 11.7, 6.7, and 1.7 nM, within a single experiment that consumes as little as 10 μl of lysates and similar volumes of the energy solution. From these reagents, we produce approximately 100 drops for each DNA concentration. We repeat the same experiment using a solution containing 7.5 nM DNA to screen additional four DNA concentrations. The resulting double emulsions are incubated for 3 h at 29°C and their intensity is quantified using fluorescent microscopy. The amount of protein produced in a drop increases with increasing amounts of DNA up to a concentration of 6 nM and levels off thereafter, as shown by black circles in Figure 4. A similar trend is seen in the experiments performed in bulk, as shown by red triangles in Figure 4. However, using drops, we obtain a 100 times improved statistics while consuming 8-fold less reagents than if the screening is performed in bulk. These results demonstrate the potential of our device to screen different reaction conditions with very low volumes of reagents.

**Figure 4:**
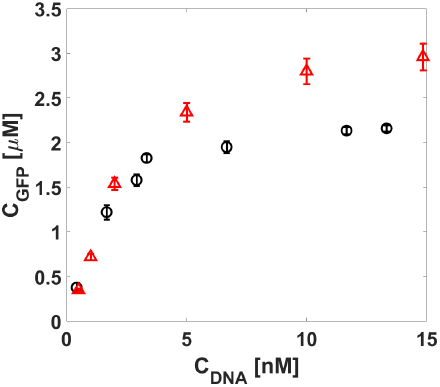
*In situ* DNA-titration. The influence of the DNA concentration on the amount of GFP synthesized inside double emulsions (○) and in bulk (Ƶ). For GFP synthesized in double emulsions, the DNA concentration was varied *in situ* by pulse width modulation.

## Conclusions

We present a microfluidic drop maker that allows mixing of up to three liquids on chip using pulsed width modulation before the resulting mixture is encapsulated in double emulsions that display a narrow size distribution. Employing green fluorescent protein synthesized in a cell-free reaction as a model system, we demonstrate that this device can be used to screen influences of the synthesis conditions on the amount of protein produced while consuming only small volumes of reagents. In addition, this device requires minimal manual operation, thereby eliminating the risk for pipetting errors. These features are of particular importance for the production of expensive biomolecules and for screening and characterization of samples that are only available in very small quantities. Hence, this device might open up new possibilities to screen synthesis conditions also for reactions that involve expensive or rare reagents.

## Conflicts of interest

There are no conflicts to declare.

## Acknowledgements

J.-C. Chang was financially supported by the EPFL Food and Nutrition center and Z. Swank by the European Research Council (ERC) under the European Union’s Horizon 2020 research and innovation program (Grant Agreement No. 723106).

## Supporting Information

### Materials and Methods

#### Materials

GFP is synthesized by mixing an aqueous solution containing cell-free reagents and an energy solution. The cell-free reagent solution contains, lysate extracted from *E. coli* and GFP DNA templates. The energy solution is composed of water containing 10.5 mM magnesium glutamate, 100 mM potassium glutamate, 0.25 mM dithiothreitol (DTT), 1.5 mM of each amino acid except leucine, 1.25 mM leucine, 50 mM HEPES, 1.5 mM adenosine triphosphate (ATP), and guanosine-5’-triphosphate (GTP), 0.9 mM cytidine triphosphate (CTP) and uridine triphosphate (UTP), 0.2 mg/mL tRNA, 0.26 mM coenzyme A (CoA), 0.33 mM nicotinamide adenine dinucleotide (NAD), 0.75 mM cyclic adenosine monophosphate (cAMP), 0.068 mM colonic acid, 1 mM spermidine, 2% PEG-8000, 4 mM 3-Phosphoglyceric acid (3-PGA).

#### Fabrication of Microfluidic Device

The microfluidic device is made of poly(dimethylsiloxane) (PDMS) using soft lithography.^1, 2^ It contains five inlets, one for the outer phase, one for the middle phase, and three for the inner phases. In addition, it contains five inlets for the control valves where air is injected to close the pneumatic valves located on top of the respective fluid channels. The maters used for the bottom part of the device that contains the liquid channels is fabricated from two layers of negative photoresist, SU-8; the first layer is 14-20 μm tall, the second layer is 100 μm tall. The masters employed to fabricate the top part of the device is made of three layers of photoresist: The first layer is 14 μm tall and composed of a positive photoresist, AZ9260, the second and third layers are 20 μm tall and composed of a negative photoresist, SU-8.

The microfluidic device is made from Sylgard 184 PDMS (Dow Corning). The three parts are joined through reactive bonding: To fabricate the top part of the device, we employ a base: crosslinker ratio of 1: 5, the middle part is made at a ratio base: crosslinker ratio of 1: 20, and the bottom part at a ratio base: crosslinker ratio of 1: 10.^3^ The middle part must be thin to ensure that the valves are sufficiently flexible to close the fluid channels if the control channels are pressurized. To control the thickness of the middle part of the device, we spin coat PDMS to form a 100 μm thick layer. PDMS is cured at 80°C for 25 minutes. The top and middle parts are aligned and bonded by incubating them at 80°C for 2 hours. The resulting part is removed from the mold and bonded to the bottom part using oxygen plasma followed by incubation at 65⁰C for 12 hours. The resulting devices have 100 μm tall control channels. The inlet liquid channels are 20 μm tall and lead into the three dimensional junction where the outermost liquid phase meets the main channel; at this junction the channel height is increased to 60 μm.

To produce water-oil-water double emulsions, the part upstream the junction where the outermost phase flows into the main channel must be hydrophobic whereas the main channel further downstream must be hydrophilic. To render the top part of the device hydrophobic we inject fluorinated oil (Novec 7500, 3M, MN) containing 1 vol% trichloro1H,1H,2H,2H-perfluorooctyl)silane (Sigma-Aldrich, MO) into this section of the device. To render the remaining part of the device hydrophilic, the surfaces are treated with an aqueous solution containing 2 wt% poly(diallyldimethylammonium chloride) and 1 M sodium chloride.

Fluids are injected into the device through polyethylene tubing (PE/5, Scientific Commodities Inc., AZ) using syringe pumps.

#### Encapsulation of cell-free reagents

We employ an aqueous solution containing 10 wt% of poly(vinyl alcohol) (PVA), *M_w_* 13 000 - 23 000 Da, 87-89% hydrolyzed, as an outer phase, a perfluorinated oil, HFE7500 (Novec 7500, 3M, MN), containing 1 wt% of surfactant^4, 5^ as a middle phase, and an aqueous phase as an inner phase. To initialize the device, we use deionized water as an inner phase. Once the device runs stably, we close the valve F that controls the fluid flow of the deionized water. Simultaneously, the two other valves, D and E, are alternatingly opened to inject the aqueous solutions containing the reagents from the two other inlets for the inner phase. Within one experiment, we inject 10 μl of an aqueous solution containing lysate extract solution through inlet D and 10 μl of an aqueous solution containing DNA and energy source through inlet E. Thereby, we keep the duty cycle of each valve at 50 ms, resulting in a total cycle time at 100 ms.

#### DNA titration

To perform DNA titration experiments, we employ three different aqueous phases as inner phases: Inlet E contains lysate with an energy source, inlet D contains an aqueous solution with 15 nM of DNA, and through inlet C we inject pure water. We change the DNA concentration, without changing the concentration of any other reagent, by varying the duty cycles of the valve G and H that control the flow of the aqueous solutions containing pure water and DNA respectively. The duty cycles are 400 ms. To enlarge the range of DNA concentrations that are screened, we repeat the same experiment but inject an aqueous solution containing 7.5 nM of DNA through inlet D. Drops are subsequently incubated at 29 °C for 3 h. During this incubation, we monitor the formation of green fluorescence protein using fluorescence microscopy where one image is acquired every 8 minutes.

#### Analysis of the formation of green fluorescence proteins

The formation of GFP is quantified using fluorescence micrographs. The average fluorescence intensity of each drop is quantified and normalized for its size and shape. To correct for lensing effects that occur at the drop interfaces, we perform control experiments where the fluorescence intensity of solutions containing known amounts of fluorescein is measured in bulk, in single emulsion, and double emulsion drops. Lensing effects increase the fluorescence intensity in single emulsions by 11%, compared to the bulk and in double emulsions by 15%. We correct for these lensing effects and convert the fluorescence intensity into a protein concentration using a calibration curve measured in bulk.

#### Drop sorting

To separate reagent-containing drops from empty ones, we introduce a T-junction downstream the drop generation junction. By pressurizing the left control channel of the sorting unit, the left valve partially closes the channel such that its hydrodynamic resistance increases and drops flow into the right outlet. To switch the direction of the fluid flow, we close the right valve 10 ms before we open the left one. If the fluid flow upstream the T-junction does not exceed 600 μl/h, drops follow the fluid flow and remain intact even when the fluid flow switches direction, as shown in the time lapse optical micrographs in Figure S1a. By contrast, if the injection flow rate exceeds 600 μl/h, some drops split at the T-junction while the direction of the fluid flow is changed, resulting in some loss of reagents, as shown in the time lapse optical micrographs in Figure S1b.

**Figure S1:**
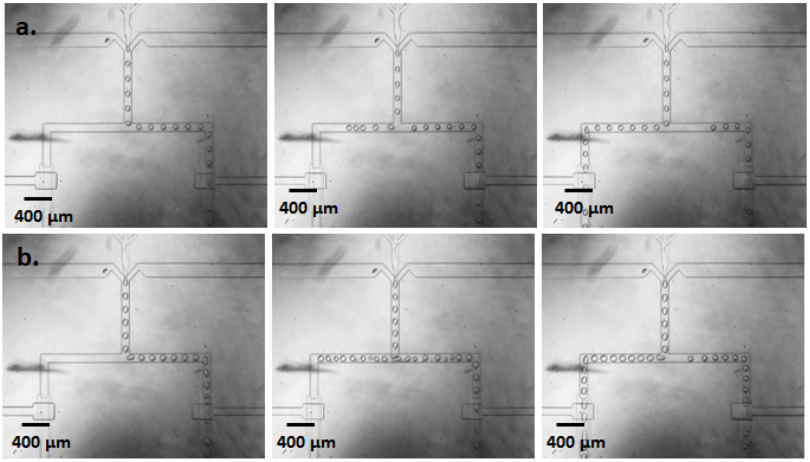
Drop sorting. Time lapse optical microscopy images illustrating the change in the flow direction of drops if the total injection speed is (a) 600 μl/h and (b) 900 μl/h.

#### Synthesis of green fluorescence proteins in drops

To test if we can track the formation of GFP inside drops, we form monodisperse single emulsion drops with a diameter of 70 μm that are loaded with lysates and 4 mM of 3-PGA. Even though the PGA concentration in these drops is almost an order of magnitude below the PGA concentration typically used, most of the drops coalesce, as indicated by the high polydispersity of drops incubated at 29°C for 30 min, shown in Figure S2a and the even higher polydispersity of drops after they have been incubated at this temperature for 3 h, as shown in Figure S2b. Coalescence of drops hampers their use for screening assays. We expect the close proximity of a high concentration of ions close to the liquid-liquid interface to deter drop stability. To test this expectation, we produce double emulsion drops containing lysates and 4 mM 3-PGA in their core; the outer liquid-liquid interface of these drops is separated from ions by their oil shell. Indeed, double emulsions are much more stable against coalescence, as indicated by their narrow size distribution after they have been incubated at 29°C for 30 min, as shown in Figure S2c and 3 h, as shown in Figure S2d.

**Figure S2:**
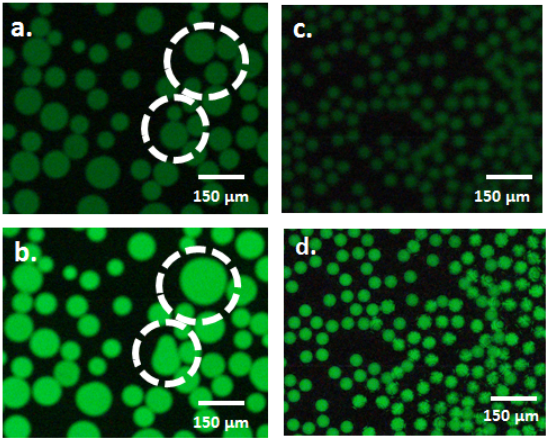
Synthesis of GFP in drops with 4 mM 3-PGA. (a-d) Fluorescence micrographs of (a, b) single emulsion and (c, d) double emulsion drops loaded with lysates an incubated at 29°C for (a, c) 30 min and (b, d) 3 h.

#### Quantification of green fluorescent protein concentrations

To quantify the amount of GFP produced in single and double emulsion drops we measure a calibration curve in bulk. The amount of GFP produced in solutions containing 4 mM 3-PGA is approximately 50% lower compared to solutions containing 30 mM, as summarized in Figure S3. By contrast, the kinetics of the GFP production is not affected by the concentration of 3-PGA, as shown in Figure 3c in the main paper.

**Figure S3:**
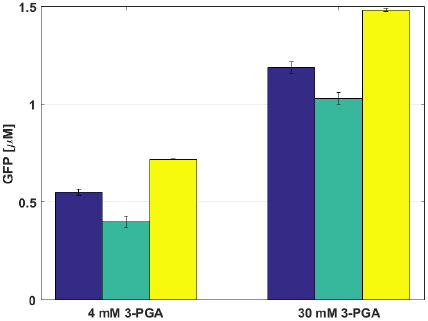
Concentration of GFP synthesized in 65 μm drops containing 4 mM and 30 mM 3-PGA. GFP is synthesized in single emulsions (blue), in double emulsions (green), and in bulk solution (yellow)

